# Time series of chicken stool metagenomics and egg metabolomics in changing production systems

**DOI:** 10.1101/2023.03.06.531264

**Authors:** Michael E. G. Rosch, Jacqueline Rehner, Georges P. Schmartz, Sascha K. Manier, Uta Becker, Rolf Müller, Markus R. Meyer, Andreas Keller, Sören L. Becker, Verena Keller

## Abstract

Different production systems of livestock animals influence various factors, including the gut microbiota. We investigated whether changing the conditions from barns to free-range impacts the microbiome over the course of three weeks. We compared the stool microbiota of chicken from industrial barns after introducing them either in community or separately to a free-range environment. Over the six time points, 12 taxa - mostly lactobacilli - changed significantly. As expected, the former barn chicken cohort carries more resistances to common antibiotics. These, however, remained positive over the observed period. At the end of the study, we collected eggs and compared metabolomic profiles of the egg white and yolk to profiles of eggs from commercial suppliers. Here, we observed significant differences between commercial and fresh collected eggs as well as differences between the former barn chicken and free-range chicken.

Our data suggest that the gut microbiota can change over time following a change in production systems. This change also influences the metabolites in the eggs. We understand the study as a proof-of-concept that justifies larger scale observations with more individual chicken and longer observation periods.

## INTRODUCTION

Over the past 50 years, the world’s population has been increasing exponentially, currently counting approximately 8 billion individuals. Researchers estimate to reach 10 billion people in 2050^1^, which in addition to rising income and global urbanisation leads to an increasing demand for food sources, in particular animal-derived foods. Poultry meat displays the fastest expanding meat production globally, with an average annual growth rate of 5 % over the past 50 years, followed by 3.1 % for pork, and 1.5 % for beef ^2^ Furthermore, egg production has been constantly increasing globally in the last decades^3^. To achieve high production efficiency, the chicken industry used to supply chicken with sub-therapeutic doses of antibiotics, provoking antimicrobial resistances in microorganisms inhabiting them. In 2003, Europe banned the preventive use of antibiotics, which resulted in an increase of systemic infections in livestock chicken, requiring the use of therapeutic doses of antibiotics^4^. The massive use of antibiotics led to an emergence of antimicrobial resistant bacteria found on poultry meat, which raises health concerns for humans^5^. Of note, bacteria themselves as natural producers present in soil or other samples are a relevant source for new sustainable antibiotics^6^.

Based on the technological advances in next generation sequencing and the gaining importance of the microbiome regarding various factors, researchers make use of genome wide shotgun sequencing to explore the microbiome in human health^7,8^. Different specimen types, extraction kits and sequencing approaches affect the results of respective studies, calling for standardized approaches^9^. With maturing technology and algorithms, researchers have also initiated projects focusing on impacts on the intestinal microbial composition of chicken and other livestock, such as diet, supplementation and living conditions^10-13^. It has been shown that the gastrointestinal tract impacts animal productivity and health^14^. In dairy cows, for example, a certain, yet dynamic gut bacteria composition was associated with greater milk production and better overall health. In chicken, a free-range housing environment positively affects egg quality of laying hens^15^. Further, the intestinal microbial composition of chicken influences eggshell quality and safety regarding offspring and human consumers^16^.

On the one hand, threats arising from caged poultry production, as well as the increasing importance of high-quality food, consumers demand free-range egg and meat production^17^. On the other hand, free-range poultry farming is more costly requiring larger surfaces and reducing sustainability. Accordingly, the exploration of hybrid keeping methods appears worthwhile.

Therefore, in this study, we aim to explore the impact of changing production systems on the microbiome and egg composition in chicken. In detail, we wanted to understand initial differences in gut microbiota composition associated with production systems and we wanted to see if a short intervention on such can lead to an approximation to the free-range chicken gut. Further, we wanted to test if, given enough time eggs appear indistinguishable among cohorts. We performed a longitudinal analysis of the intestinal microbiome of free-range chicken, compared to previous barn chicken which were transferred to a free-range environment using fecal samples over the course of 8 weeks. Additionally, we investigated metabolites in the egg white and egg yolk of store-bought eggs of barn and free-range chicken and compared the metabolite composition to the chicken we kept in a free-range environment, those that were placed into a free-range environment coming from a barn industry, and chicken that originated from a barn industry and were put into a free-range community of existing free-range chicken.

## RESULTS

We defined a study set-up that lets us conclude on the change of the stool microbiomes and get additional information on the eggs as secondary read out. We acquired six barn chicken, which were primarily held for industry purposes and randomly split them into two cohorts of three chicken (Fig. 1a). The first cohort consisting of chicken H1-H3 remained isolated with access to a small chicken coop with an enclosure. The second cohort, H4-H6, was released from the industry barn and joined another cohort of existing free-range chicken, H7-H8, in a different coop with enclosure. For each chicken, we monitored microbial composition in their feces at timepoints day 0, 3, 7, 10, 14, and 21 and performed genome-wide metagenomic sequencing. Of note, the collection approach enabled us to unambiguously match the stool samples to the individual chicken because each sample was picked immediately. Of note, the gut microbiome of chicken is known to depend on the collection site within the digestive system. We, however, collected excreted samples. Especially the feces originating from the comparably large ceca and non-ceca parts (Fig. 1b) of the gut differ, calling for attention during the data analysis. Even though visual differences exist, molecular profiling facilitates additional insights on the origin of the sample picked.

**Figure 1:**
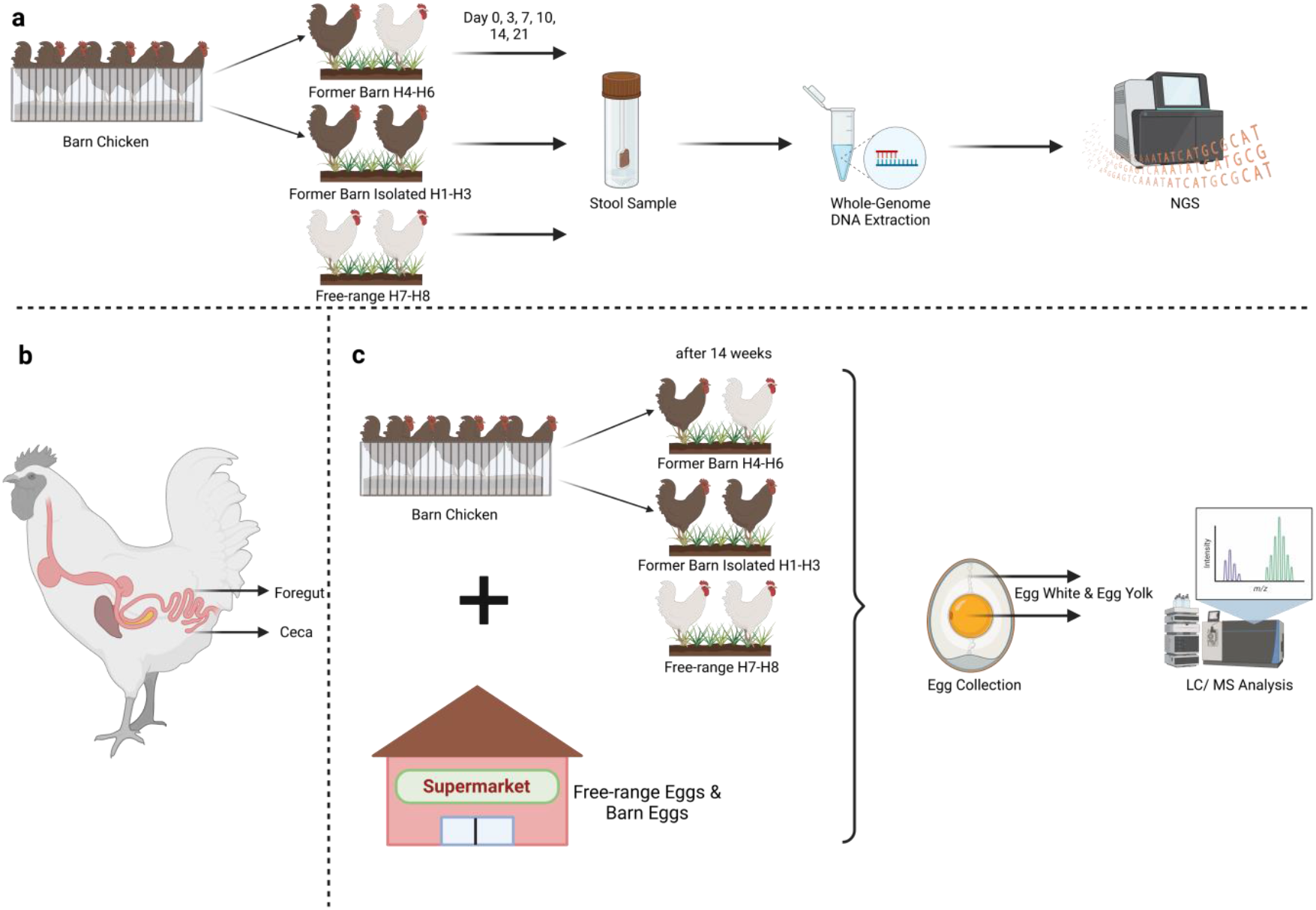
Study design and sample collection. **a)** Cohort design, fecal sample collection and subsequent analyses. Barn chicken (depicted as brown chicken) were released from an industry barn and divided into two cohorts. H1-H3 were released into a free range, isolated from existing free-range chicken (depicted in white). H4-H6 were released and grouped together with existing free-range chicken. Samples were also collected from long-term existing free-range chicken H7 and H8. Fecal samples were collected from all cohorts on several days and subjected to wholegenome DNA extraction and subsequent whole-genome sequencing. **b)** From all cohorts, eggs were collected 97 days past study begin and the egg white and egg yolk of each egg were analysed by mass spectrometry. Results were compared to eggs and eggs from barn chicken from a local supermarket. Additionally, an egg from a newly relocated former barn chicken (entered the environment the day of egg collection), which was transferred into a free-range environment and grouped with H7 and H8 was analyzed.

Additionally, to the fecal samples. we also collected eggs of the various cohorts 97 days after study start for the duration of 7 days. We then analyzed the metabolite composition of the egg white and egg yolk of each egg by mass spectroscopic analyses. Further, we compared the results with the metabolite composition of free-range eggs and barn chicken eggs bought in a local supermarket (Fig. 1c).

### Fecal microbiome composition varies upon release

Following sequencing we performed a stringent quality control (QC) including removal of host DNA from the metagenomic samples and other QC steps (c.f. methods). The QC identified on average 0.57% (1.9% ± SD) of reads to be of low quality per sample. Afterwards, we performed taxonomic profiling where we observed an estimated average 50.18% (16.5% ± SD) assignment rate on a metagenomic assembled genome/strain level. With these taxonomic profiles, we initiated a first ordination analysis. Here, we observed no patterns linked to either time or the cohort (Fig. 2a). We thus repeated a similar analysis with MinHash similarities where we again did not observe any clusters or explainable trends associated with the two features (time and cohort) of interest (Fig. 2b). While these analyses suggest no general trend with respect to the cohort and time, the spread of the points motivates a closer look. Focusing on the relative microbiome compositions, we observed major variations and heterogeneity across samples (Fig. 2c). One group of samples (including amongst others H2 day 0, H3 day 7, or any of the samples by H7) are mostly composed of *Lactobacillus, Limsilactobacillus*, and *Ligilactobacillus*. Another group (including most measurements, e.g., H4 day 21, H5 day 10 and H8 day 10) differs significantly. The samples of the second group are characterized by a larger diversity on the genus level. The first set of samples falls in line with the foregut chicken microbiome that is extensively discussed in literature^18,19^. The larger diversity in the composition of the second group can be attributed to ceca samples in chicken, as discussed above (compare to Fig. 1b). Accordingly, we repeated the previous MinHash similarity analysis but extended our samples with the data generated by Huang et al.^20^, providing an atlas of chicken metagenomes in different gut regions. Indeed, we confirm the expected clustering: most samples with lower *Lacto-, Limsilacto-*, and *Ligilactobacillus* relative abundances clustered closer to the cecum samples, whereas other samples scattered into the ileum, duodenum, and jejunum region (Fig. 2d). As expected, the major variations within the samples are mostly due to the detection of either the foregut or hindgut microbiome. To minimize effects associated with gut regions in further analyses, we decided to subset our data to retain only the samples that clustered into the foregut region of the previous analysis. On the remaining 29 samples, we performed differential abundance analysis at the species level and performed three different statistical tests, comparing i) differences in the microbiome due to initial production systems, ii) changes in the microbiome in relocated chicken over time iii) differences in changes over time comparing the two resettled cohorts. Combining all three test results, a total of twelve different OTUs showed statistical significance (adjusted alpha level of 0.01) (Fig. 2e). Four of the twelve species were significantly differentially abundant in all three tests (Fig. 2f).

**Figure 2:**
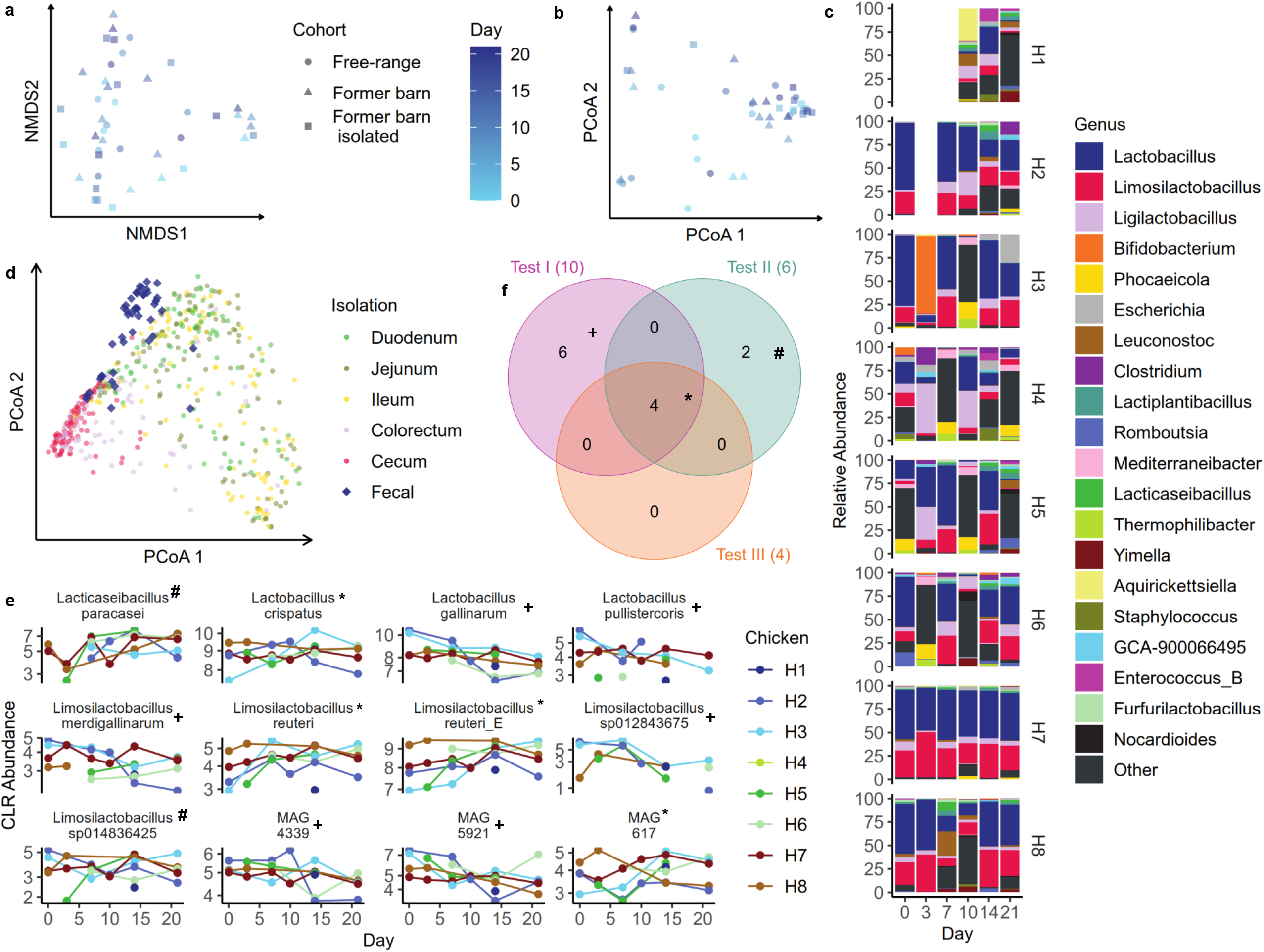
Stool Microbiota Composition. **a)** Non-metric Multidimensional Scaling (NMDS) of Bray-Curtis dissimilarities computed on species information of gathered metagenomic samples after quality controlled. Time of sampling is indicated by color, whereas point shaped designates the cohort. **b)** Principal coordinate Analysis computed on 1-MinHash similarity. Point shape shows sampling cohort, colors represent sampling time after study kick-off. **c)** Relative species abundance on genus level for the different timepoints and chickens. **d)** PCoA of Mash dissimilarities computed on our own samples extended by the dataset by Huang et al. All highlighted fecal samples are from this study. **e)** Operational taxonomic units that were highlighted as significant after p-value adjustment in at least one of the three statistical tests. The different intersections (compare to panel f are marked by +, # and *, respectively) **f)** Visualized concordance among test results. Each set denotes one statistical test and entries indicate the number of statistically significant differentially abundant OTUs.

### Increased number of resistances seems to remain in free-range environment

Among the most relevant aspects in livestock farming are resistances against antimicrobials^21-25^. We thus searched for potential antimicrobial resistance mechanisms across all metagenomic samples. Consistently over all cohorts, we identified microbial resistance against tetracyclines, which is an antibiotic frequently used in veterinarian medicine. Further, macrolide resistance appears to be prevalent in most chicken. In consideration of the One Health aspect, vancomycin resistance detected in all cohorts is a considerable threat, as bacteria carrying the resistance might also transfer onto humans and add onto the global health crisis of antimicrobial resistances. Vancomycin is commonly used as a last resort drug as resistance development is usually slow^26^. Moreover, we found bacteria resistant to carbapenems in the cohort, which was released from the industry barn and placed into an isolated free-range environment. Carbapenem resistance in Gramnegative bacteria is mainly caused by carbapenemases and counts as a major and on-going global health problem which is spreading rapidly and causing serious outbreaks with limited treatment options (Fig. 3a). Interestingly, the carbapenem resistance found in the industry barn chicken is caused by the subclass B1 metallo-beta-lactamase JOHN-1, which was previously described in *Flavobacterium johnsoniae* but not detected in human colonizing bacteria yet^27^. A relevant aspect is the number of present resistances in the different cohorts. We thus computed the average number of resistances for the original barn chicken and the free-range chicken at the beginning of the observation period and at the end after three weeks (Fig. 3b). For both time points, we recognize an increased number of resistances for the industrial barn chicken. This number seems however not to change over time, i.e., present resistances remain in the stool microbiota for at least three weeks. Comparing the early and late time point average resistances (Fig. 3 c) and those in the free-range and former barn cohort separately (Fig. 3d) emphasizes this trend. Of note, we only reach statistically significant differences if all early and late time points are pooled together (Wilcoxon Mann-Whitney p-value 0.016) due to the limited number of chickens in the cohorts. Having identified changes in microbiota over time and having observed resistance factors in the metagenomes, we asked on differences in the eggs.

**Figure 3:**
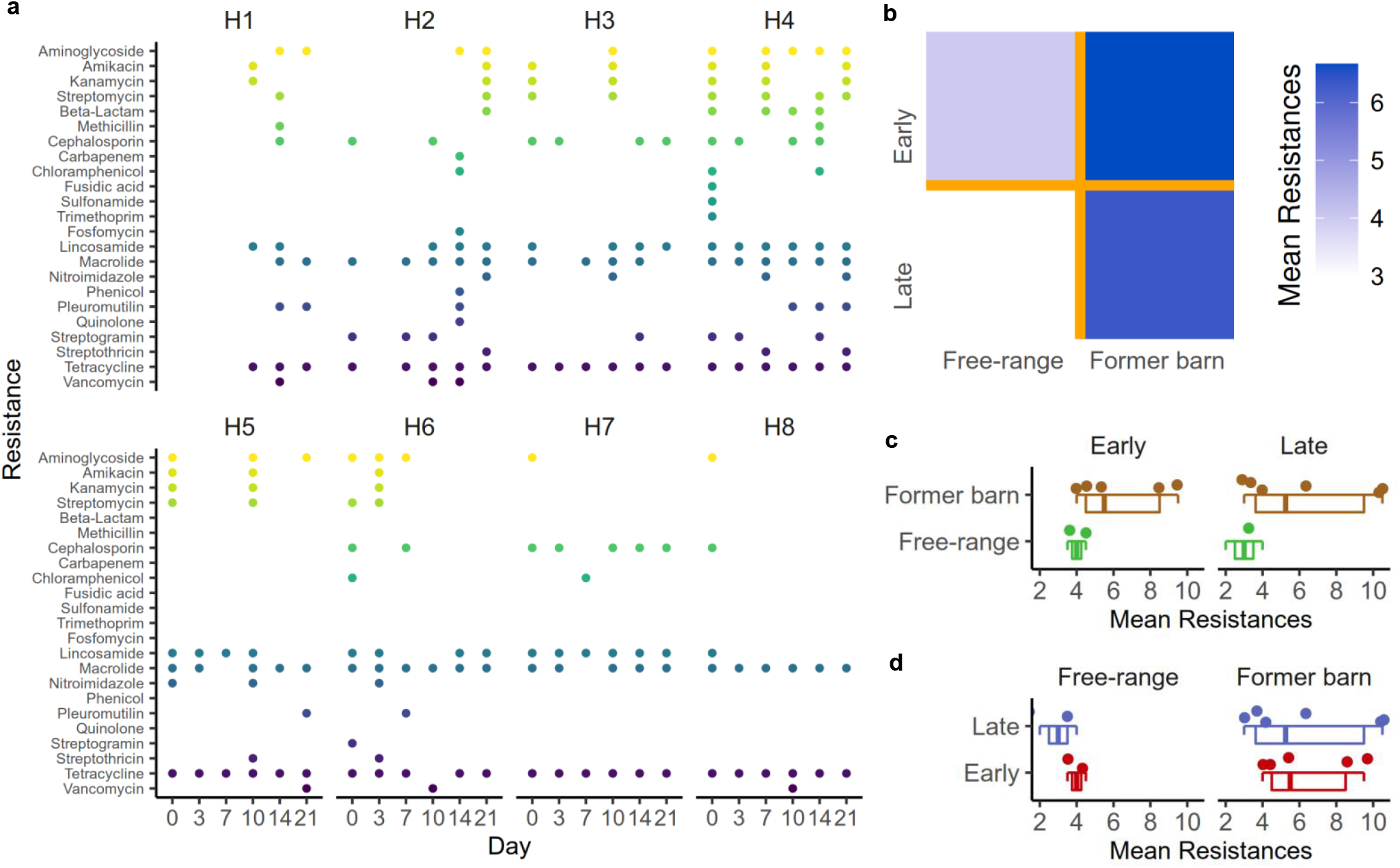
Resistances in free-range and caged chicken. **a)** Detected resistances with ABRicate in the different assembled metagenomic samples. **b)** Boxplots with individual data points showing the averaged resistances in the early and late time point split by the cohorts. **c)** Boxplots with individual data points showing the averaged resistances in the cohorts split by the early and late time points.

### Metabolomic profiles of egg differ between free-range and barn as well as commercial eggs

To assess differences in egg composition we performed untargeted HPLC-MS of egg yolk and egg white separately. After peak calling and bias correction, we searched two different sets of features. First, we aimed to find features that separate our collected eggs by initial production system. Second, we aimed to identify features that separate shop eggs (organic eggs from a supermarket) from fresh eggs. Initially, we selected only significant ANOVA features. For the first test, we found no feature to be statistically significant after p-value adjustment. In the second test, we observed 14 and 90 features in egg white and egg yolk respectively to meet our criteria. The small number of features undercutting the significance threshold of 0.05, the high dimensional feature spaces, and the low expected assignment rate in targeted MS, did not allow us to limit our further analysis only on ANOVA significant features. Instead, we decided to select high variance features allowing for reasonable separation after a first filtering based on ANOVA analysis. Naturally, principal component analysis (PCA) on these ANOVA pre-filtered features already displayed clear separation for our cohorts, as well as separation among bought and collected eggs (Fig. 4a). Based on the most important features in the high variance principal components of the PCA, we trained linear discriminant analyses and attained an average cross-validation accuracy of 96% (6.2% ± SD) (Fig. 4b). Asking about the nature of the identified features, we performed a targeted MS approach. We successfully identified 10 of the most relevant features identified in the PCA via targeted MS (Fig. 4c). Breaking them up in the different classes, six features derived from lipids, three from amino acids and one was matched to a flavonoid. Post hoc testing identified in 6 of the 10 cases the eggs from the supermarket to be the most significantly differing cohort. In four cases, however, the egg from originally free-range chicken differed from all other eggs, including the former ban chicken that were put in the free-range setting.

**Figure 4:**
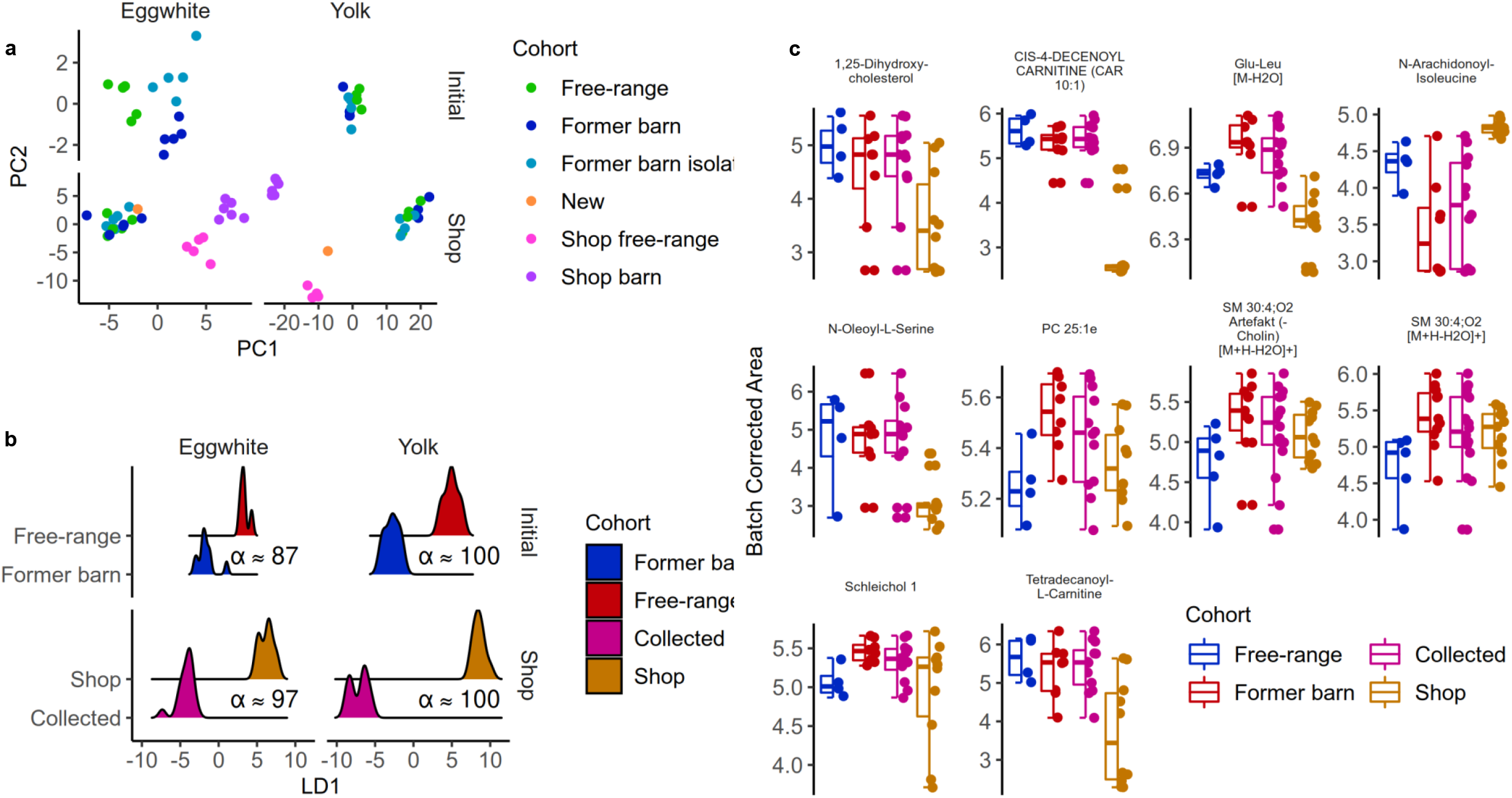
Metabolomics profiles of eggs. **a)** PCA of batch-corrected areas on ANOVA prefiltered features derived from untargeted HPLC-MS of eggs. **b)** Projections of the four different trained LDA classifiers. Visualization includes the training data. **c)** Batch-corrected peak areas of driving features we were able to identify with targeted MS/MS.

## DISCUSSION

Our study analysed the fecal microbial composition over time of two different chicken cohorts being introduced to a free-range environment, isolated and within a community of existing free-range chicken, and a control cohort. We aimed to identify potential differences in initial gut microbiota composition of free-range compared to barn chicken and detect changes over time under the different conditions mentioned above. Further, we investigated metabolite composition of egg yolk and egg white for each cohort and drew the comparison to store-bought free-range and barn eggs.

The applied statistical analysis highlighted 12 different species to have differing abundances among initial production systems or over time. *Lacticaseibacillus paracasei* was initially higher in existing free-range chicken (H7, H8) and increased over time in both released cohorts. *L. paracasei* is believed to have regulatory effects on chicken gut immunity, microbial composition, and an overall higher microbial diversity as its presence was correlated with the enrichment of the genera *Anaerotignum, Coprococcus, and Massilimicrobiota*^*28*^. *Anaerotignum* sp. produce so called short-chain fatty acids^29^, which play a key role in health homeostasis and inflammation reduction. *Corprococcus* sp. were correlated with the presence of other bacteria, which have a positive effect on host health ^28^. Therefore, an enrichment in *L. paracasei* seems favourable for chicken. H3, which was released into a group of existing free-range chicken further displays an increase in *Lactobacillus crispatus*, which belongs to the most prevalent species in the chicken gastrointestinal tract and ^30^ its presence is correlated with a protection against infectious diseases in poultry^31,32^. We further observed an initially lower abundance of *Lactobacillus gallinarium* in the existing free-range chicken (H7, H8), and a decrease of *L. gallinarium* in those chicken, which were released and grouped with H7 and H8. *L. gallinarium* is currently not correlated with any substantial positive or negative effect on poultry health or fitness^32^. However, as a lactic acid producing bacterium, *Lactobacillus* sp. are overall associated with health homeostasis and many species belonging to this genus display well-known probiotics in the animal industry^33^. Last, we observed an increase in *Limosilactobacillus reuteri* (former *Lactobacillus reuteri)* in all former barn chicken and an initially higher abundance of *L. reuteri* in the free-range chicken. In new-born chicken, supplementation with *L. reuteri* has been shown to suppress growth of non-beneficial Proteobacteria, while it promoted enrichment of rather beneficial *Lactobacillus sp*^*3*^. Also in humans, *L. reuteri* displays enormous potential as a probiotic strain^35^. As of today, little to nothing is known about *Lactobacillus pullistercoris, Limosilactobacillus merdigallinarum, Limosilactobacillus sp012843675* and *sp014836425, MAG4339 (Ligilactobacillus), MAG5921 (Lactobacillus)*, and *MAG617 (Limosilactobacillus)*, which were highlighted in our analysis (Fig. 2e). Overall, we conclude that chicken in a free-range environment show higher abundances of beneficial bacteria, and that resettling chicken from an industry barn to a free-range environment has a positive effect on health-associated bacterial abundances. Nevertheless, we want to underline that statistical analysis may be impacted by the initial sampling strategy. When combining our analyses with existing microbiome data on different gastrointestinal parts of chicken, it is obvious that our first group clusters with samples extracted from the cecum, and our second group clusters separated from all other parts of the chicken gastrointestinal tract (Fig. 2d). Previously, it could be shown that the microbial composition in the small intestine differs from those of the relatively large cecum of chicken, the large intestine, and the colorectum^20^. Chicken are capable of releasing cecal content towards the ileum or towards the cloaca, then called the cecal drop^36^. This can occur every 8-10 hours. Based on the results depicted in Fig. 2b, c, and d, we believe that some of the collected fecal samples belong to the group of cecal drops and rather represent the cecum microbial composition, whereas all other samples represent a mixture of all parts of the gastrointestinal tract, which is released with normal feces release. Depending on the selected separation criteria of these clusters, differential abundance analysis may highlight other species.

Concerning the metabolomic egg composition ANOVA results indicated reliable differences between bought and collected eggs. On the contrary, statistical analysis did not highlight differences in our collected eggs comparing initial production systems. This may likely be due to a shrinkage in overall sample size. When we looked at overall variances in a total of four investigations where we did observe variations in corrected peak area. However, whether these differences hold in the general case, would require a larger study cohort. On the one hand, our measurements of e.g., Schleichol 1, which belongs to the flavonoids, or PC25:1^37^ which are generally positively connotated in literature were non-significantly reduced in our free-ranged cohort. On the other hand, in the same cohort, we observed e.g., 1,25-Dihydroxy-cholesterol^38^ to be slightly increased, which is associated with cell homeostasis and a precursor for steroid hormones. Accordingly, we cannot clearly advocate for a production system if we use egg composition as only quality criterion.

Our study set up that includes the immediate collection of stool samples and allowing an analysis on the individual chicken also leads to challenges. First, the cohort sizes are limited because only a small number of chickens can be overseen. Here, camera-based systems with automated image analysis might support to handle larger cohorts. Further, even using molecular metagenomic measurements, the distinction between microbiota from the foregut and the cecum is challenging. The nucleic acid based read out bears further challenges: the observed resistances are encoded in the genome and do not reflect actual resistances. Additionally, the number of time points and the follow up period of three weeks may not be sufficient and an adaption of former ban chicken to the free-range chicken may happen. We thus plan to re-analyze stool microbiota of the cohort after a one-year interval. Further, the data suggest other downstream analyses such as the identification of novel bioactive gene clusters that might be a source of novel antibiotics.

We thus consider our study as a proof-of-concept justifying further analyses at larger scale. Using time series and individually matched metagenomes bears an enormous potential to provide further insights in the dynamics of livestock metagenomes. Finally, high-resolution metagenomics might enable an improved surveillance of arising resistances in animal populations.

## METHODS

### Study design and sample collection

We acquired six chickens from a local industry barn in July 2022 and divided them into two cohorts (n=3). H1-H3 were released from the industry barn and transferred to a free range but were kept isolated from other free-range chicken with access to a small chicken coop with an enclosure, and ii) H4-H6 released from the industry barn and joined an existing cohort of free-range chicken (H7, H8). From all eight chickens, we collected fecal samples on day 0, 3, 7, 10, 14, and 21. All fecal samples were subjected to whole-genome DNA extraction and subsequent whole-genome sequencing. Further, after 97 days past study begin, we collected eggs from all study cohorts (H7, H8 n=5, H1-H3 n=5, H4-H6 n=5) and analyzed the metabolite composition of egg white and egg yolk via mass spectrometry. We compared the results with the analyses of store-bought free-range eggs (n=5) and eggs from barn chicken (n=6). We further analyzed an egg from a newly resettled former barn chicken, which was transferred directly into an existing herd of free-range chicken. The respective egg was collected on the day of relocation (Fig. 1).

### DNA extraction

DNA was extracted from all fecal samples using the ZymoBIOMICS DNA Miniprep Kit. The DNA was extracted according to the manufacturer’s protocol. Briefly, 50 mg of fecal samples were used for DNA extraction according to the manufacturer’s recommendation. The mechanical lysis of bacterial cells was performed using the MP Biomedicals™ FastPrep-24™ 5G Instrument (Fisher Scientific GmbH, Schwerte, Germany). The velocity and duration were adjusted to 6 m/s for 45 s three times with 30 s of storage on ice in between each lysis step. DNA was eluted in 20 μl DNase/RNase free water^9^. The DNA concentration was determined via NanoDrop 2000/2000c (Thermo Fisher Scientific, Wilmington, DE) full-spectrum microvolume UV-Vis measurements.

### Library Preparation and sequencing

Extracted whole-genome DNA was sent to Novogene Company Limited (Cambridge, UK) for library preparation and sequencing. Briefly, samples were subjected to metagenomic library preparation and further sequenced via paired-end Illumina Sequencing PE150 (HiSeq). For all samples, 5 Gb reads per sample were generated.

### Sequencing data analysis

As the first step of metagenomics shotgun sequencing analysis, we cleaned the reads by removing host sequences and performed QC with BBduk (version (v):38.98, command line arguments (cla): “k=23 mink=11 hdist=1 ktrim=r tbo tpe out=stdout.fq -ref illumina.fasta” & “maq=10minlength=50k=31 mcf=0.5 -ref GRCg6a.fasta”) and GRCg6a as a reference. QC information was visualized with MultiQC (v1.11)^39^. For MinHash based comparison and taxonomic profiling we computed signatures on cleaned reads with sourmash^40^ (v4.4.3, cla:”sketch dna -p k=21,k=31,k=51,scaled=1000,abund --merge”). Comparison between samples was based on k-mer size 31. Own reanalysis of samples from Huang et al.^20^ followed the same data cleaning and signature computation pipeline. To prepare taxonomic profiling, we extended the Genome Taxonomy Database (GTDB ; vGTDB R07-RS207)^41^ with metagenomic assembled genomes from Segura-Wang et al.^11^ and Feng et al.^42^. To this end MAGs of both studies were dereplicated together with GTDB using drep^43^ (v:3.4.0 cla “comp 50 -con 10-- checkM_method lineage_wf --S_algorithm fastANI --S_ani 0.95 -nc 0.5”). All dereplicated MAGs that were not part of GTDB were retained and their signatures and indexes were computed with sourmash. Taxonomic profiling was performed with sourmash on quality-controlled reads with k-mer size 51 using the union of GTDB and the previously dereplicated MAGs as a reference. All further downstream analysis of taxonomic profiles was performed in R relying on the phyloseq package^44^ (v1.40.0). Ordination analysis was computed with non-metric multidimensional scaling on Bray-Curtis distances. For all three differential abundance analysis with ANCOMBC^45^ (v:1.6.2) we kept only the samples that resembled a foregut signature as described in the results section. Further, for every test, we set the significance level at 0.01, the prevalence cutoff at 0.8, and the p-value adjustment method to “Benjamini-Hochberg”. For the first test comparing initial conditions we subset the remaining samples to only keep samples from day 0 and compared cohort of existing, free-ranged chicken with those of former barn chicken. In the second and third test we used all samples except those derived from the control group. The second test performed regression on the number of days, whereas the last test looked at the interaction term of days and final production systems. Metagenomic assembly was performed with spades (v:3.15.4, cla: “--meta”). Resistance annotation was done with ABRicate (v:1.0.1) and the NCBI AMRFinderPlus database (v)

### Metabolomics sample preparation

All eggs were separated and egg white and yolk were frozen for one week at -20 °C. Afterwards, 400 μL methanol:ethanol (1:1, v/v) containing the internal standards 48 μM tryptophan-d5, 53.4 μM glucose-d_7_, 34.8 palmitic acid-d31, and 8.6 μM creatinine-d_3_ were added to 100 μL egg white or 100 mg egg yolk. All samples were shaken for 2 min at 2000 rpm and subsequently centrifuged for 30 min at 15,000 rpm and 2 °C. 150 μL of the supernatant were transferred into a new reaction tube and evaporated to dryness using a vacuum centrifuge (Concentrator plus, Eppendorf, Hamburg) at room temperature and program “V-aq” for roughly 4 h. The residue was reconstituted in 100 μL of a mixture containing methanol and acetonitrile (30:70, v/v) by shaking for 5 min at 15,000 rpm and 22 °C. Ten μL of each sample of both drugs of abuse were pooled to obtain one quality control sample (group QC) for every cell experiment. Every obtained sample was transferred into an amber glass vial and 1 μL was injected onto the HPLC-HRMS/MS as described below.

### LC-HRMS/MS apparatus for metabolomics

The analysis was performed using a Thermo Fisher Scientific (TF, Dreieich, Germany) Dionex UltiMate 3000 RS pump consisting of a degasser, a quaternary pump, and an UltiMate Autosampler, coupled to a TF Q Exactive Plus system equipped with a heated electrospray ionization HESI-II source. Mass calibration was done prior to analysis according to the manufacturer’s recommendations using external mass calibration. Additionally, before each experiment, the spray shield and capillary were cleaned. The performance of the column and mass spectrometer was tested using a mixture described by Maurer et al. before every experiment. The conditions were set according to published procedures^46,47^. Gradient reversed-phase elution was performed on a TF Accucore Phenyl-Hexyl column (100 mm x 2.1 mm, 2.6 μm, TF, Dreieich, Germany) or on a hydrophilic interaction liquid chromatography (HILIC) Nucleodur column (125 × 3 mm, 3 μm, Macherey-Nagel, Düren, Germany) for normal-phase chromatography. The mobile phases for gradient elution using the Phenyl-Hexyl column consisted of 2 mM aqueous ammonium formate containing acetonitrile (1 %, v/v) and formic acid (0.1 %, v/v, pH 3, eluent A), as well as 2 mM ammonium formate in acetonitrile and methanol (1:1, v/v), containing water (1 %, v/v), and formic acid (0.1 %, v/v, eluent B). The flow rate was set from 1-10 min to 500 μL/min and from 10-13.5 min to 800 μL/min using the following gradient: 0-1.0 min hold 99 % A, 1-10 min to 1 % A, 10-11.5 min hold 1 % A, 11.5-13.5 min hold 99 % A. Normal-phase chromatography was performed using aqueous ammonium acetate solution (200 mM, eluent C) and acetonitrile containing formic acid (0.1 %, v/v, eluent D). The flow rate was set to 500 μL/min using the following gradient: 0-1 min 2 % C, 1-5 min to 20 % C, 5-8.5 min to 60 % C, 8.5-10 min hold 60 % C, 10-12 min hold 2 % C. For preparation and cleaning of the injection system, a mixture containing isopropanol and water (90:10, v/v) was used. The following settings were used: wash volume, 100 μL; wash speed, 4000 nL/s; loop wash factor, 2. Every analysis was performed at 40 °C column temperature, maintained by a Dionex UltiMate 3000 RS analytical column heater. The injection volume for metabolomics analyses was 1 μL and for those analyses investigating the formation of imines 10 μL. The HESI-II source conditions for every experiment were as follows: ionization mode, positive or negative; sheath gas, 60 AU; auxiliary gas, 10 AU; sweep gas, 3 AU; spray voltage, 3.50 kV in positive mode and -4.0 kV in negative mode; heater temperature, 320 °C; ion transfer capillary temperature, 320 °C; and S-lens RF level, 50.0. Mass spectrometry for UM was performed according to a previously optimized workflow^48^ using full scan (FS) only. The settings for FS data acquisition were as follows: resolution, 140,000 at *m/z* 200; microscans, 1; automatic gain control (AGC) target, 5 × 10^5^; maximum injection time, 200 ms; scan range, *m/z* 50-750; polarity, negative or positive; spectrum data type, centroid.

Settings for parallel reaction monitoring (PRM) data acquisition were as follows: resolution, 35,000 at *m/z* 200; microscans, 1; AGC target, 5 × 10^5^; maximum injection time, 200 ms; isolation window, 1.0 *m/z;* collision energy (CE), 10, 20, or 40; spectrum data type, centroid. The inclusion list contained the monoisotopic masses of all significant features, and a time window of their retention time 30 s. Analysis was performed using a randomized sequence order with five injections of pure methanol (reversed-phase chromatography) or eluent D (normal-phase chromatography) samples at the beginning of the sequence for apparatus equilibration, followed by five injections of the pooled QC sample. Additionally, one QC injection was performed every five samples to monitor batch effects as described by Wehrens et al.^49^. TF Xcalibur software version 3.0.63 was used for all data handling.

### Metabolomic Data analysis

Thermo Fisher LC-HRMS/MS RAW files were converted to mzML format using Proteo Wizard^50^ and subsequently parsed by XCMS^51^ in an R environment for raw data inspection and peak picking. Total ion chromatograms, base peak chromatograms and mean intensity chromatograms were visually inspected for deviations that may hint to improper measurements. Additionally, total ion currents of the samples were monitored in boxplots to discover batch effects during measurements. The quality of the peak picking and the alignment was monitored using total ion chromatograms and extracted ion chromatograms of the used internal standards. Annotation of isotopes, adducts, and artifacts was performed using the package CAMERA^52^. Optimization of XCMS parameters was in accordance with a previously optimized strategy. Peak picking and alignment parameters can be found in the according Jupyter notebooks. Names of the features were adopted from XCMS using “M” followed by the rounded mass and “T” followed by the retention time in seconds (e.g., “M218T222” for an ion at *m/z* 218.1538 and a retention time of 222 s). Before log transformation, missing values were replaced by the lowest measured peak area as proposed by Wehrens et al.^49^ as a surrogate limit of detection. Batch correction was performed for each HPLC experiment by performing linear regression on QC samples, predicting each peak area based on the position of the investigated sample in the respective experiment. Based on these regression models, areas of all peaks for all samples were corrected by subtracting residuals and adding the average peak area. After area correction, we aggregated all peaks across all yolk 4 HPLC experiments into one feature vector and repeated the same for egg white. We removed QC samples for all further analysis. We then proceeded to analyze both egg components independently. Further, we made minor changes to the following workflow as needed depending on the hypothesis we tested, resulting in 4 workflows. For the first hypothesis comparing only collected eggs from our chicken, we removed the egg from the newest chicken for further analysis. The second hypothesis comparing shopped eggs included this egg. For both hypotheses we performed ANOVA on both egg components with Benjamini-Hochberg correction *as /*-value adjustment method. For yolk feature filtering was applied on unadjusted ANOVA p-value threshold of 0.01. For egg white we used 0.05. On the remaining features and datapoints, PCA was performed. We then computed feature loadings on principal components with a higher-than-average variance, with a minimum of three first principal components. The highest absolute loadings were selected for targeted MS analysis. We performed linear discriminant analysis only on high variance principal components using Monte-Carlo cross-validation.

### Identification of significant features

MS^2^ spectra were recorded using the above mentioned PRM method to allow identification of significant features. Individual spectra were exported after subtracting the baseline left and right of the peak. After conversion to mzXML format using Proteo Wizard, spectra were imported to NIST MSSEARCH version 2.3. A library search for identification was conducted using the following settings: Spectrum Search type, Identity (MS/MS); Precursor Ion m/z, in spectrum; Spectrum Search Options, none; Presearch, Off; Other Options, none. MS/MS search was conducted using the following settings: Precursor tolerance, ± 5 ppm; Product ion tolerance, ± 10 ppm; Ignoring peaks around precursor, ± m/z 1. The search was conducted by using the following libraries: NIST 14 (nist_msms and nist_msms2 sublibraries), Wiley METLIN Mass Spectral Database, HMDB 5^53^ (MS/MS Experimental), LipidBlast^54^, and MassBank (NIST). Additionally, LipidMaps COMP_DB search^55^, MetFrag^56^, and MONA similarity search were used for inconclusive spectra matches or those that were not matched in MSSEARCH. Metabolites of the investigated NPS were identified by comparing and interpreting their spectra to those of the parent compounds.

## Acknowledgements

We want to acknowledgement DFG for supporting this by providing us with a Q Exactive Plus.

## Author Contributions

MEGR, UB, AK, RM and VK had the idea for this study, together with GPS, SKM and JR had full access to all the data and take responsibility for the integrity of the data and the accuracy of data analysis. MEGR and UB collected samples. JR and SLB extracted whole-genome DNA. Computational data analysis was performed by GPS and AK. JR, GPS, SKM, AK and VK drafted the manuscript. All authors critically reviewed the paper for important intellectual content and agreed to submit the final version for publication.

## Data Availability

Metagenomic and metabolomic data will be made available upon publication. Reviewer link will be made available upon request.

